# Minilungs from hESCs to study the interaction of *Streptococcus pneumoniae* with the respiratory tract

**DOI:** 10.1101/2022.02.01.478764

**Authors:** Fernando González-Camacho, Suélen Andreia Rossi, Julio Sempere, María Pilar de Lucas, José María Rojas-Cabañeros, Carlos Pelleschi Taborda, Óscar Zaragoza, José Yuste, Alberto Zambrano

**Affiliations:** Biotechnology of stem cells and organoids, Chronic Diseases Program, Institute of Health Carlos III, Madrid, Spain; Spanish Pneumococcal Reference Laboratory, Spanish National Microbiology Centre, Institute of Health Carlos III, Madrid, Spain; Cellular Biology Unit, Chronic Diseases Program and CIBERONC, Institute of Health Carlos III, Madrid, Spain; Department of Microbiology, Biomedical Sciences Institute, University of São Paulo (USP), São Paulo, Brazil; Mycology Reference Laboratory, Spanish National Microbiology Centre, Institute of Health Carlos III, Madrid, Spain; CIBER of Respiratory Diseases (CIBERES)

**Keywords:** Minilungs, human pluripotent stem cells, human embryonic stem cells, hESCs, *Streptococcus pneumoniae*, *pneumococcus*, surfactant proteins, alveolar cells, disease modeling

## Abstract

The new generation of organoids derived from human pluripotent stem cells holds a promising strategy for modeling host-bacteria interaction studies. Organoids recapitulate the composition, diversity of cell types and, to some extent, the functional features of the native organ. We have generated lung bud organoids derived from human embryonic stem cells to study the interaction of *Streptococcus pneumoniae* (pneumococcus) with the alveolar epithelium. Invasive pneumococcal disease is an important health problem that may occur as a result of the spread of pneumococcus from the lower respiratory tract to sterile sites. We show here an efficient experimental approach to model the main events of the pneumococcal infection that occur in the human lung exploring bacterial adherence to the epithelium, internalization, and triggering of an innate response that includes the interaction with the surfactant and the expression of representative cytokines and chemokines. Thus, this model, based on human minilungs, can be used to study pneumococcal virulence factors, the pathogenesis of different serotypes and it will allow therapeutic interventions in a reliable human context.

**IMPORTANCE:** *Streptococcus pneumoniae* is responsible for high morbidity and mortalities rates worldwide affecting mainly children and adults older than 65 years. Pneumococcus is also the most common etiologic agent of bacterial pneumonia, non-epidemic meningitis, and a frequent cause of bacterial sepsis. Although the advent of pneumococcal vaccines has decreased the burden of the diseases caused by pneumococcus, the emerging of antibiotic-resistant strains and non-vaccine types by serotype replacement, is worrisome. To study the biology of pneumococcus and to establish a reliable human model for pneumococcal pathogenesis, we have generated human minilungs from embryonic stem cells. The results show that these organoids can be used to model some events occurring during the interaction of pneumococcus with the lung such as adherence, internalization, and the initial alveolar innate response. This model also represents a great alternative to study virulence factors involved in pneumonia, drug screening, and other therapeutic interventions.

## OBSERVATION

*Streptococcus pneumoniae* (pneumococcus) is the leading cause of severe bacterial pneumonia with high morbidity and fatality rates worldwide. Lower respiratory infections were responsible for over 2.74 million deaths in 2015, and the third major cause of mortality worldwide in children under 5 years of age, being *S. pneumoniae* the leading cause of these infections (1). There are currently two vaccines to prevent Invasive Pneumococcal Disease (IPD), the 13-valent conjugate vaccine (PCV13) that can be used for children and adults, and the 23-valent polysaccharide vaccine only indicated for adults. During the last decade, the interest to characterize the complex interplay between host-microbes interactions involving organoid cultures has increased.

Organoids share important features with the original organ that make them attractive models to study the pathogenesis or the microbial colonization of the tissue. For instance, the variety of cell types, the spatial arrangement of cells, and some level of organ functionality. It has been reported a plethora of examples of organoids derived from adult stem cells and pluripotent stem cells (PSCs) to study the interaction microbe-epithelium and pathogenesis. Hence, organoids from the human liver, stomach, intestine/colon or gall bladder, have been used to explore infections of protozoan parasites and bacteria such as *Escherichia coli, Helicobacter pylori, Clostridioides difficile* and *Salmonella typhimurium* (2). The natural biology of pneumococcal interactions at the respiratory tract includes the nasopharyngeal colonization, production of acute otitis media, sinusitis and, in many cases, the dissemination to the lower respiratory tract producing pneumonia. The pneumococcal carrier state is usually an asymptomatic event for the vast majority of individuals and it is a prerequisite for infection of the lower respiratory tract, producing pneumonia, bacteremia and, eventually, meningitis. During the early stages of pneumonia, pneumococcus avoids the mucociliary and antibacterial peptide barriers by the expression of different virulence factors (3). Adhesion to lung cells involves interactions with host cell glycoconjugates followed by interaction with host cell protein receptors that promote internalization (4). As the alveolar epithelium is denuded, the underlying extracellular matrix of bronchioles and alveoli are exposed. Host-pneumococci interactions in the alveoli result in inflammation and cytotoxicity. At the point where alveoli are overfilled with bacteria, edema fluids, erythrocytes and fibrins, bacteremia could eventually take place. Finally, dissemination to the bloodstream may occur by damaging the epithelium and direct invasion of endothelial cells. While the knowledge about these processes mentioned above is abundant, little is known about the specific interaction of pneumococcus with alveolar type I and II (ATI and ATII) cells (responsible for gas exchange and surfactant production) or their responses during the initial infection or the early phases of acute pneumonia. This is partially due to the lack of reliable cell models and the difficulty to extract, establish, and maintain, primary ATI and ATII cells in culture. At the alveoli, ATII cells represent the preferred target for pneumococcal adherence and cell damage (5) (6) (7) (8). An important defense system of the lung is the surfactant that reduces the surface tension, preventing, therefore, alveolar collapse at the end of expiration and protecting the lung from injuries and infections. The surfactant is a mixture of phospholipids, proteins and carbohydrates, which is produced and secreted by the ATII cells. Surfactant protein A (SFTPA) has an opsonizing activity that mediates phagocytosis and removal of several respiratory microorganisms including *S. pneumoniae* (9) (10). SFTPB peptides seem to mediate aggregation of both Gram-positive and Gram-negative bacteria and their elimination by permeabilization of the bacteria cell membrane. As SFTPA, SFTPD binds polysaccharides located on the bacterial surface and promotes aggregation, phagocytosis, complement activation, and inhibition of bacterial colonization and invasion (11) (12) (13) (14) (15).

We report here the characterization of a human organoid model suitable for studying the pneumococcal interactions with the human alveolar epithelium. We used human embryonic stem cells (hESCs; AND-2 line) to build lung bud organoids (LBOs) that develop branching airway and early alveolar structures embedded in Matrigel™ sandwiches. As previously shown, the expression pattern and structural features indicated that those LBOs structures may reach the second trimester of human gestation (16). Lung organoids, as the generated here, have numerous advantages over cell lines or simple primary cultures as they offer an unlimited availability of primary cells and can emulate structural and functional features of the original organ. The generation of optimal lung organoids relies primarily on the selection of good hESCs showing adequate morphology when growing along with inactivated feeder cells [inactivated mouse embryo fibroblasts (iMEFs)]. hESCs are picked-up and passaged to new plates with iMEFs to accumulate material for the subsequent differentiation to lung lineage. Figure 1A shows the aspect of a good hESC colony, the expression of pluripotency markers (NANOG and SOX2), and representative micrographs at various times of the differentiation process: Embryoid bodies (EBs), anterior foregut endoderm (AFE) and cultures at day 17 representing nascent organoids in suspension. Figure 1B shows the expression of lung field specification markers NKX2.1 and FOXA2 (at differentiation day 22) by indirect immunofluorescence and RT-qPCR. As previously reported, these two markers show a pan-nuclear homogeneous expression in nearly the whole cell population indicating the correct differentiation to the lung field. Figure 1C shows the microscopic aspect at day 56 of LBOs embedded in Matrigel™ sandwiches undergoing the adequate state of differentiation characterized by the presence of lung buds more or less branched as previously described (16) (17). Figure 1C also shows a RT-qPCR result illustrating the complexity of these cultures that express high levels of alveolar epithelial cell markers: PDPN (ATI cells) and the surfactants (ATII cells). As previously described by our group and others (16) (17) (18) (19) (20), the differentiation protocol applied here yields cultures enriched in alveolar epithelial cells although specific markers of other cells (airway epithelial cells) could also be found in proximal locations and after long-term time maintenance of these structures (16). Figure 1D shows a representative micrographs of a histochemical analysis (H&E staining) denoting the typical branches of the LBOs at day 60. We performed microinjections of LBOs branches at different locations to assure the optimal delivery of pneumococci to the lumen as we previously described for recombinant adenovirus (21). Further characterization of the LBOs generated included the expression of the surfactant proteins and the proper visualization of pneumococci by confocal microscopy (Supplemental Figure 2). The expression of the surfactant proteins illustrates the maturity level of the organoid and the bias to ATII cells, a preeminent target of pneumococcus at the alveoli. Pneumococcal cells were easily observed in the lumen of the branches, attached at the cell surface, or immersed at the cytoplasm of the cells delineating the distal branches of the organoid. The analysis by confocal microscopy yielded the unequivocal interaction between pneumococcus with surfactant proteins (Supplemental Figure 2). To model the pneumococcal infection within the alveolar epithelium, we delivered pneumococci into the lumen of the organoids and follow the interaction, in this case, with SFTPD, at different times. Microinjected organoids were processed for immunofluorescence, and colony counts were quantified at different post-infection times (2, 4, 8, and 24 h). Figure 2A illustrates the transit of the bacteria through the organoid epithelium demonstrating the proper delivery of the bacteria to the lumen, adherence to the ATII cell surface, and internalization. At t0 there were multiple pneumococci at the lumen (green staining, panel t0). At 4 and 8 h, we observed the presence of numerous pneumococcal cells in the lumen (green staining; green arrowheads), interacting with SFTPD at different locations and infecting the cytoplasm of ATII cells (merged staining; yellow arrowheads). The panel corresponding to 24 h of Figure 2A shows the significant decrease in the number of pneumococci (green staining) inside the ATII cells and its absence at the LBO lumen, indicating, very likely, the clearance of the bacteria. In addition, we observed the presence of few but particularly large pneumococci-SFTPD aggregates inside the cell [merge staining; yellow arrowheads Figure 2A, at 24 h]. The dynamics delineated by this microscopic analysis fit well with the numbers of colony-forming units (CFUs) found in extracts of the microinjected organoids at different times post-infection. Hence, at t0 (organoids processed immediately after the microinjection), CFUs in the order of 10^8^ were be found in the cell extracts (Figure 2B). A fast decay in the numbers of viable bacteria was observed between 2 h and 4 h and, from 4 h to 24 h. The lack of viable bacterial counts in the extracts is related to the absence of pneumococcal cells in the immunofluorescence (green staining) and the presence of those large pneumococcus-SFTPD aggregates observed at 24 h (Figure 2A). The early interaction of the surfactant with the bacteria described here correlated with the innate response of alveolar cells to the pneumococcal infection. ATII cells and, to a lesser extent ATI cells, are essential effector cells in the inflammatory response to pneumococcal infection even when they cover around 5% of the alveolus surface (22). Upon infection, ATII cells can release a plethora of antimicrobial molecules, cytokines and chemokines (22)(23). These mediators contribute to the migration of monocytes and macrophages to the site of infection and their activation. Figure 2C shows the RT-qPCR result corresponding to different components of the innate response induced by the organoids infected with pneumococcus. We analyzed *TLR2, IL6, IL8, TNFα, CXCL5* and *CCL20* (*MIP3*) because they are important components of the host immune response involved in the recognition and the inflammatory response to invading pathogens such as *S. pneumonia*e (7) (8) (23) (24) (25). Our results indicate a rapid response of the alveolar epithelium to the exposition of pneumococci. We also observed that pneumococcal infection of the lung organoids triggered the expression of many of these mediators reaching a peak at 4 h, and *TLR2* showing an upward trend of expression. This is consistent with previous data showing the relevance of TLR2 which is expressed on the surface of ATII cells, and it has been shown to allow NF-κB activation after pneumococcal infection (26).

**Figure 1.**
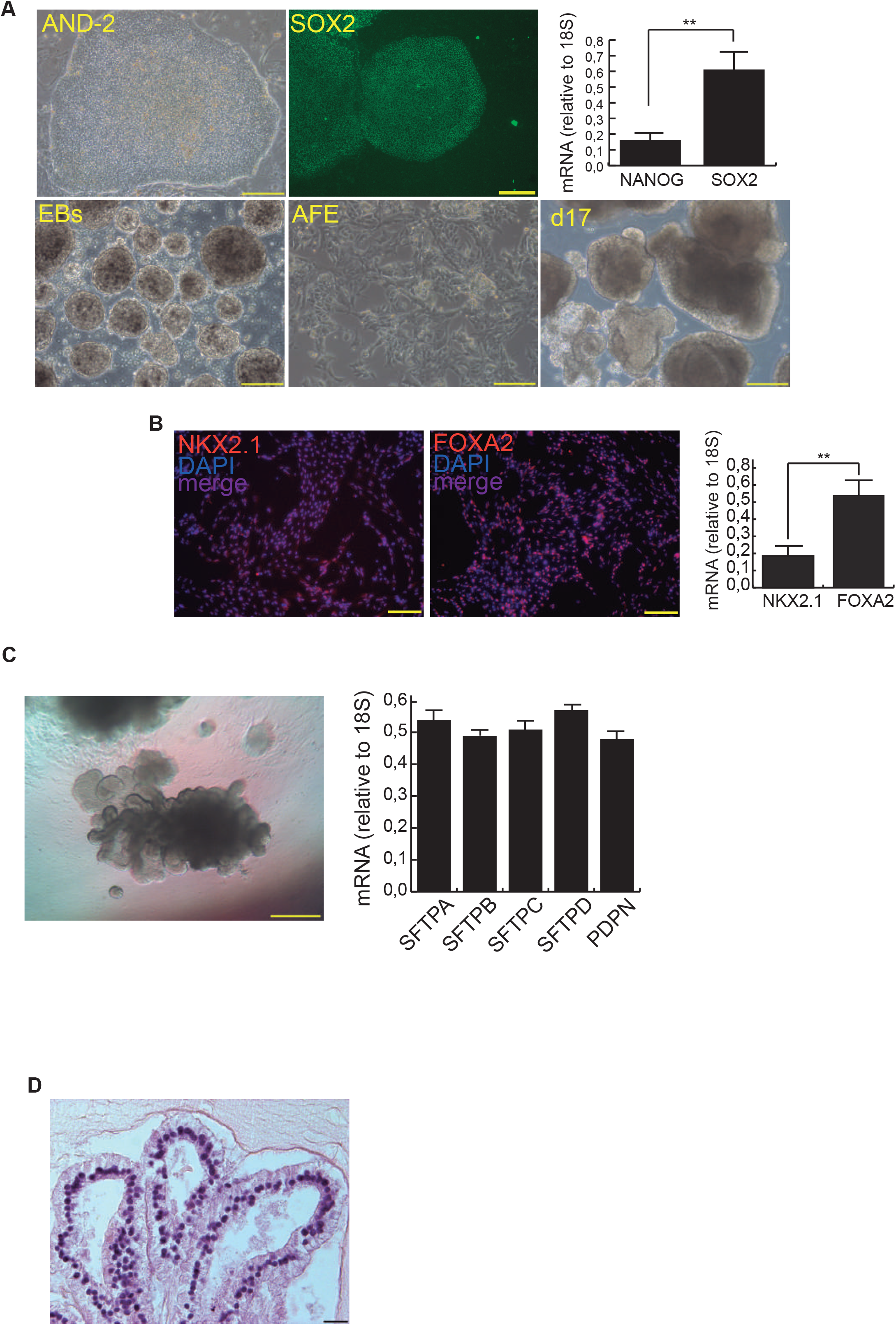
Generation of human minilungs: Sequential differentiation process and expression markers. **A**. Left upper micrograph: AND-2 colony growing along with feeder cells [inactivated MEFs (iMEFs)]; scale bar: 100 μm; central upper micrograph: Expression of SOX2 in an undifferentiated colony of AND-2; scale bar: 100 μm. Right upper panel: RT-qPCR analysis of pluripotency markers *NANOG* and *SOX2* (n=3; >4 organoids per experiment were used; P=0.0029). Bottom panels: Representative micrograph of EBs (embryoid bodies), AFE (anterior foregut endoderm) and representative micrographs of cultures at day 17 of differentiation (nascent organoids) scale bar: 100 µm. **B**. Detection of FOXA2 and NKX2-1 (markers of the lung field) by indirect immunofluorescence and RT-qPCR results at day 22 (n=3; > 4 organoids per experiment were used; P=0.0027). Negative controls of these immunofluorescences can be found in the Supplemental Figure 1. **C**. Representative micrograph of lung bud organoids embedded in Matrigel™ sandwiches and expression levels (relative to 18S) of alveolar epithelial cells markers (day 56) [(n=3; >4 organoids per experiment were used; ANOVA P=0.0674]; scale bar: 100 µm. **D**. Histochemical analysis of LBO sections (H&E staining) of organoids at day 60; scale bar: 50 µm.

**Figure 2.**
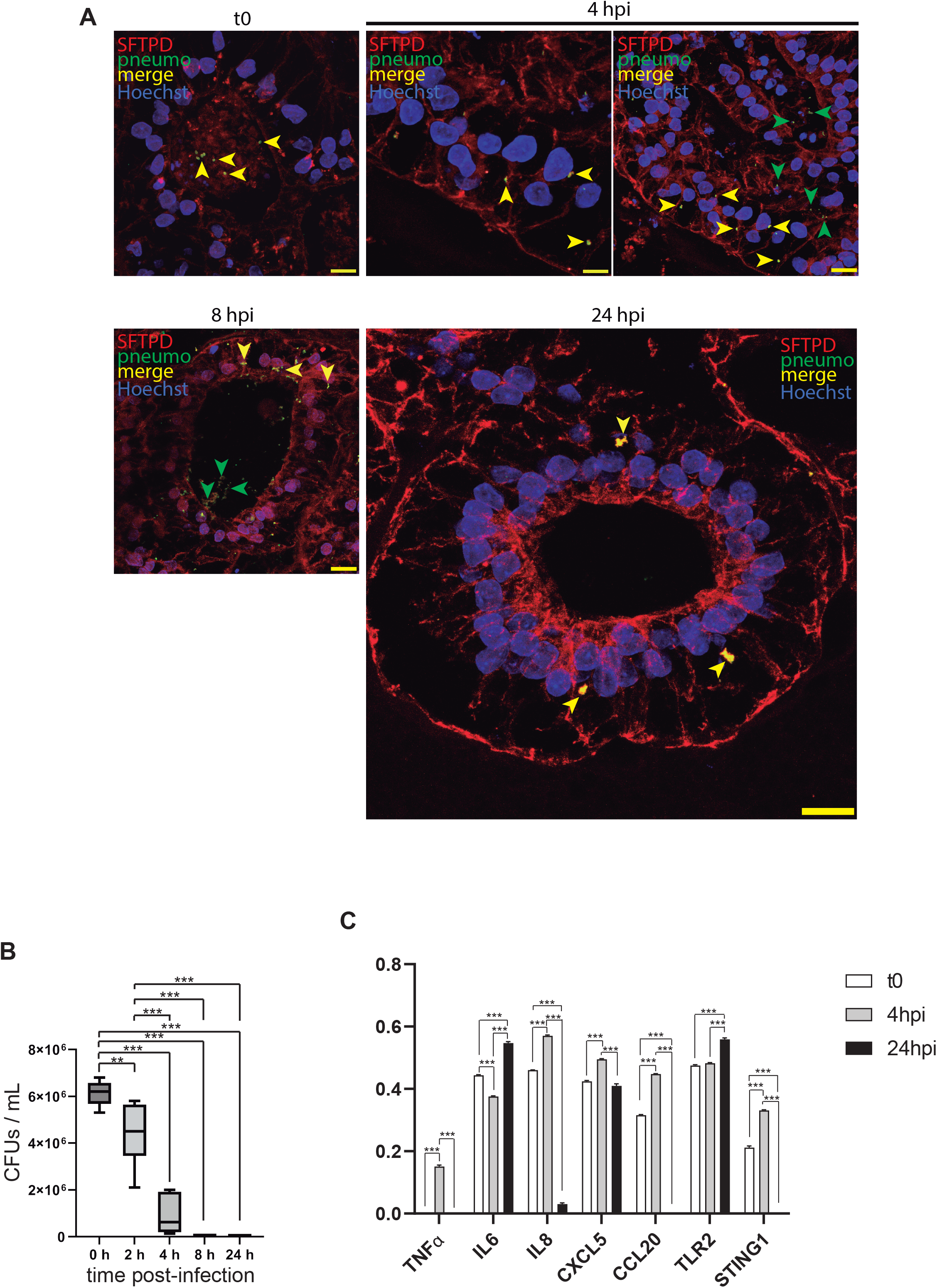
Interaction of pneumococcus with the surfactant system of the microinjected organoids. **A**. Interaction of pneumococcus with lung buds at different times post-microinjection (t0 to 24 hpi); t0: Arrowheads signal pneumococcus particles at the lumen of the organoid; 4 hpi: Green arrowheads and yellow arrowheads signal pneumococcus particles at the lumen of the organoid and inside the alveolar epithelium (invaded epithelium), respectively; 8 hpi: Green arrowheads and yellow arrowheads signal pneumococcus aggregates at the lumen of the organoid and inside the alveolar epithelium, respectively; 24 hpi: Yellow arrowheads signal pneumococcus and SFTPD aggregates inside the alveolar epithelium. **B**. Left panel: Quantification of CFUs/ml [n=3; >4 organoids per condition were used; ANOVA P<0.0001]. **C**. RT-qPCR result of markers representing the alveolar innate response to pneumococcus infection at different times post-infection. *IL6* (Interleukin 6), *IL8* (C-X-C motif chemokine ligand 8), *TNF*α (Tumor necrosis factor), *TLR2* (Toll-like receptor 2), *STING1* (Stimulator of interferon response cGAMP interactor 1), *CXCL5* (C-X-C motif chemokine ligand 5*), CCL20* (C-C motif chemokine ligand 20) [n=3, >4 organoids per experiment were used; ANOVA P<0.0001]. The results presented in the figures are means ± SEM. Significance of the analysis is indicated as, *: P<0.05, **: P<0.01, ***: p<0.001.

Overall, our study demonstrates the suitability of hESCs derived lung organoids to model early interactions of *S. pneumoniae* with the alveolar epithelium including the host immune response. Below we summarize the potential of the model presented here in terms of human material availability, time, and relevance to pneumococcal biology:

i. Human primary cells are organized in 3D structures
ii. Unlimited and rapid availability of precious and primary human material
iii. Infective bacteria and other targets can be delivered directly to the lumen by microinjection to mimic alveoli exposure to exogenous agents
iv. It has an enormous potential to study the adherence of bacteria to the eukaryotic cell and the influence of bacterial virulence factors. This potential includes the possibilities of therapeutic interventions: developing antiadhesive strategies to block transmission and colonization.
v. It facilitates the study of common elements in the pathogenesis of all serotypes to evaluate antigenic formulations for novel pneumococcal vaccines
vi. It will allow the establishment of specific individual organoids, organoid biobanks, and high-throughput analyses.

## SUPPLEMENTAL MATERIAL includes

- Material and Methods
- Supplemental references for material and methods
- Supplemental Figures
- Supplemental Table

## DECLARATIONS

### ETHICS APPROVAL AND CONSENT TO PARTICIPATE

The use of the hESC line AND-2 and the experimental procedures of this study were approved by the ISCIII Ethics Committee (ref. no. CEI PI 10_2015-v2) and the National Committee of Guarantees for the Use and Donation of Human Cells and Tissues (ref. no. 345 288 1 and 436 351 1).

### CONSENT FOR PUBLICATION

Not applicable

### AVAILABILITY OF DATA AND MATERIALS

Please contact the corresponding author for data requests.

### AUTHORS’ CONTRIBUTIONS

FGC and SAR performed and designed experiments, analyzed data. MPL performed experiments. JS performed experiments, analyzed data, and wrote the paper. JMRC and CPT analyzed data. OZ performed and designed experiments, analyzed data. JY designed experiments, analyzed data, and wrote the paper. AZ performed and designed experiments, analyzed data, wrote the paper, and conceived the project. The authors read and approved the final manuscript.

## ACKNOWLEDGEMENTS

We thank the histology facility (Manuel and Marta) of the ISCIII for technical help.

## LIST OF ABBREVIATIONS

AFE: Anterior Foregut Endoderm
ATI: Alveolar Type I Cells
ATII: Alveolar Type II
Cells: BMP4 Bone Morphogenic Protein 4
BSA: Bovine serum albumin
*CCL20*: C-C motif chemokine ligand 20
*CXCL5*: C-X-C motif chemokine ligand 5
EBs: Embryoid Bodies
FBS: Fetal bovine serum
FGF: Fibroblast Growth Factor
*FOXA2*: Forkhead box A2
hbFGF: Human basic fibroblast growth factor
hESCs: Human Embryonic Stem Cells
*IL6*: Interleukin 6
*IL8*: *CXC8*, C-X-C motif chemokine ligand 8
KGF: Keratinocyte growth factor
LBOs: Lung bud organoids
MEFs: Mouse Embryonic Fibroblasts
*NANOG*: Nanog homeobox
*NKX2-1*: NK2 homeobox 1
*OCT3/4A*: *POU5F1*, POU class 5 homeobox 1
PBS: Phosphate-buffered saline
*PDPN*: Podoplanin
RT-qPCR: Quantitative Real-Time RT-PCR (Reverse Transcription Polymerase Chain Reaction)
SEM: Standard error of the mean
SFD: Serum-free differentiation
*SFTPA*: Surfactant Protein A
*SFTPB*: Surfactant Protein B
*SFTPC*: Surfactant Protein C
*SFTPD*: Surfactant Protein D
*SOX2*: SRY (Sex Determining Region Y)-box 2
*STING1*: Stimulator of interferon response cGAMP interactor 1
*TLR2*: Toll-like receptor 2
*TNFα*: *TNF*, tumor necrosis factor
μm: Micrometer

